# Profiling the plasmid conjugation potential of urinary *E. coli*

**DOI:** 10.1101/2022.03.02.482680

**Authors:** Cesar Montelongo Hernandez, Catherine Putonti, Alan J. Wolfe

## Abstract

*Escherichia coli* is often associated with urinary tract infection (UTI). Antibiotic resistance in *E. coli* is an ongoing challenge in managing UTI. Extrachromosomal elements – plasmids – are vectors for clinically relevant traits, such as antibiotic resistance, with conjugation being one of the main methods for horizontal propagation of plasmids in bacterial populations. Targeting of conjugation components has been proposed as a strategy to curb spread of plasmid-borne antibiotic resistance. Understanding the types of conjugative systems present in urinary *E. coli* isolates is fundamental to assessing the viability of these strategies. In this study, we profile the two well studied conjugation systems (F-type and P-type) in the draft genomes of 65 urinary isolates of *E. coli* obtained from bladder urine of adult women with and without UTI-like symptoms. Most of these isolates contained plasmids and we found that conjugation genes are abundant/ubiquitous, diverse, and often associated with IncF plasmids. To validate conjugation of these urinary plasmids, we conjugated the plasmids from two urinary isolates, UMB1223 (predicted to have F-type genes) and UMB1284 (predicted to have P-type genes), into the K-12 *E. coli* strain MG1655. Overall, the findings of this study support the notion that care should be taken in targeting any individual component of a urinary *E. coli* isolate’s conjugation system, given the inherent mechanistic redundancy, gene diversity, and different types of conjugation systems in this population.

## Background

*E. coli* is linked to urinary tract infection (UTI)^1–3^. Because the key genetic determinants of *E. coli* pathogenicity in the urinary tract remain unknown, management is a challenge^4–6^. This clinical dilemma is compounded by antibiotic resistance^7,8^. Genes that encode antibiotic resistance are often carried by plasmids, genetic elements that can propagate and retain genes not only for resistance but also virulence and fitness^8–10^. Due to their variable genetic composition, plasmids can be classified by incompatibility (*inc*) grouping, which denotes the inability of two plasmids to be maintained in the same host^11–13^. A prime example of clinically relevant plasmids in *E. coli* are the F plasmids, characterized by the *inc* loci – *incFI* and/or *incFII*^14,15^. F plasmids are widespread, heterogenous, stable, and generally mobile^8,10,16^. F plasmids have frequently been found in urinary *E. coli,* specifically in extended spectrum β-lactamase (ESBL)-producing *E. coli* isolated from urinary infection^17^. Also, the uropathogenic *E. coli* (UPEC) strain UTI89 harbors an F plasmid hybrid – pUTI89 – and its loss results in a significant decrease in pathogenicity in a mouse model during early infection^18^. The pUTI89 plasmid has been noted to carry pathogenesis genes in addition to a conjugation system, which is key for plasmid mobility^6,16,19^.

Conjugation is the cell-to-cell translocation of genetic information assisted by conjugative machinery – conjugative pilus, DNA processing enzymes (relaxasome), mating pair stabilization proteins, and type IV secretion system (T4SS)^16,20^. The prototypical conjugation system in *E. coli* is the F-type system, which uses transfer *(tra)* genes^10,21,22^. The transfer operon is composed of genes that are considered to be essential for conjugation *(traA, traB, traK, traC, traW, trbC, traE, traL, traH, traG, traU, tra/, traM, traD)* for F plasmid transfer and, in some cases, more than a dozen accessory genes^19,20,22^. Because expressing conjugation machinery is an expensive endeavor, it is repressed by the fertility inhibition (FinOP) system that targets the transcription regulator TraJ^23^. TraJ upregulates the transfer operon, resulting in the expression of components involved in forming the relaxosome *(traD, traI, traM, traY),* conjugative pilus *(traP, traE, traL, traC, traW, trbC, traF, trbB),* mating pair stabilization *(traN, traG, traU),* and T4SS core proteins *(traB, traK, traV*)^16,20,21,23^. The transfer operon also can encode proteins that defend against excessive conjugation with the same or similar plasmids, either by surface exclusion (TraT) or entry exclusion (TraS)^23,24^. F plasmids are the typical group associated with the F-type conjugation system^16^.

Functional and structural relatives of the F-type transfer system exist, such as the P-type VirB/VirD4 system, which also utilizes a T4SS for conjugation and has been extensively studied in *Agrobacterium tumefaciens*^16,20,25^. Components consist of the type IV coupling protein (VirD4), inner membrane complex (VirB4), outer membrane complex (VirB7, VirB9, VirB10), conjugative pilus (VirB2 pilin), transglycosylase (VirB1), and pilus-tip protein (VirB5)^16,26^. The plasmids RP4, R388, and pKM101 are well studied *E. coli* plasmids associated with a P-type system^27–29^. The VirB/VirD4 system has been implicated not just in intra-species genetic exchange, but also in inter-species and even inter-kingdom conjugation^30,31^. Likely due to the constant genetic exchange in plasmids that results in genomic heterogeneity, bacteria may carry plasmids that borrow components from different conjugation systems^16,25^.

Conjugation is a potential target in curtailing antibiotic resistance spread in the clinical setting^26,32^. Hypothetically, blocking key transfer functions by conjugation inhibitors (COINs) could decrease plasmid exchange, and by extension, decrease propagation of bacterial genes for antibiotic resistance, virulence and fitness^33–37^. It has been noted that COINs should ideally not inhibit bacterial growth and be used in conjunction with antibiotics^32^. However, deployment of such a strategy requires further considerations^26,32^. Outside of identifying genes essential for conjugation and determining how to mechanistically target them, we must know how a particular bacterial population depends on a given conjugation system. Targeting a conjugation component would be less effective if this factor was not abundant in a population or if this factor could be replaced by another component. Furthermore, if there were multiple distinct conjugation systems in a population, targeting of one system might simply lead to selection and propagation of the other^38,39^. In considering conjugation as a viable target for inhibition in the clinical setting, we must first understand in more detail the conjugation systems in clinical bacterial isolates in a given niche.

In this study, we profile F-type and P-type conjugation genes in 65 urinary *E. coli* isolated from catheterized bladder urine of adult women with or without UTI-like symptoms^5^. Plasmids were present in 84.6% (N=55/65) of urinary *E. coli* isolates, and the majority included F-type conjugation genes with the minority having the Type-P conjugation genes. To validate conjugation of these urinary plasmids and viability of the F-type and P-type systems, we conjugated the plasmids from two isolates, UMB1223 which has F-type genes and UMB1284 which has P-type genes, into the *E. coli* K-12 strain MG1655. Multiple types of conjugation loci exist in urinary *E. coli.* Overall, our investigation finds that conjugation systems are common in urinary *E. coli* and that conjugation genes are diverse in number and composition. This knowledge is important to understanding conjugation in the clinical setting and how this process may be targeted.

## Methods

This study uses the urinary *E. coli* isolates, sequence reads, and contigs first published in Garretto et al. 2020 (**Supplemental Table 1**)^5^. These isolates were recovered from urine samples collected from adult women with and without UTI-like symptoms via transurethral catheterization during several Institutional Review Board-approved studies at Loyola University Chicago (LU203986, LU205650, LU206449, LU206469, LU207102, and LU207152) and the University of California San Diego (170077AW). Assembled whole genome sequences (WGS) for these isolates were downloaded from BioProject PRJNA316969.

To identify known plasmid *inc* loci, *E. coli* WGS FASTA files were scanned using PlasmidFinder v2.1 (https://cge.cbs.dtu.dk/services/PlasmidFinder/) using the Enterobacteriaceae database with a threshold of 95% identity and minimum 60% coverage (**Supplemental table 2**)^13^. The genomes were binned into a plasmid group associated with their *inc* genes via exclusion criteria: genomes with *incF* genes were placed in the IncF group, genomes with *col* but no *incF* genes were placed in the Col group, and genomes with *inc* genes that were neither *incF* nor *col inc* genes were in the Inc-various group. The *E. coli* plasmid literature was reviewed for genes used in plasmid conjugation, primarily from the F-type transfer and P-type VirB-VirD4 systems (**Supplemental Tables 3 and 4**)^20,23^. NCBI Local BLAST v2.11.0 was used to query this initial set of conjugation genes in the 65 urinary *E. coli* draft genomes^40,41^.

Conjugation was utilized as the method to transfer plasmids from urinary *E. coli* isolates to a laboratory *E. coli* K-12 strain (MG1655 with a chloramphenicol resistance cassette) that possessed a chloramphenicol selection marker^42^. To attempt conjugation, we searched for compatible antibiotic resistance traits in urinary isolates – potential conjugation donors – CamS and resistant to one of the following – Kan, Spec, Tet, Amp. Urinary *E. coli* isolates with identified plasmids, the appropriate resistance markers, and carrying transfer genes – were grown on various antibiotic plates to assess their selection marker profile. Lysogeny broth (LB) plates (1.0% bacto tryptone, 1.0% yeast extract, 0.5% NaCl, 1.5% agar) were supplemented with these respective antibiotics: ampicillin (Amp, 100 ug/ml), chloramphenicol (Cam, 25 ug/ml), kanamycin (Kan, 40 ug/ml), spectinomycin (Spec, 100 ug/ml), or tetracycline (Tet, 15 ug/ml) (**Supplemental Table 5**). Isolates that grew on an antibiotic had their sequences reviewed for antibiotic resistance genes in their contigs, which also were compared to plasmid entries in the NCBI nr/nt database. Urinary *E. coli* plasmid donor candidates were filtered on the following basis: they could grow on an antibiotic selection marker, they carried a gene that was predicted to encode antibiotic resistance, this resistance gene was on a predicted plasmid, and they had evidence of near complete F-type or P-type operons (**Supplemental Table 6**)^16^.

Two urinary *E. coli* (UMB1223, UMB1284) met the criteria and were used as plasmid donors; UMB0939 was used as a negative control, as it met all the criteria except identifiable conjugation genes. Conjugation in *E. coli* was performed as described previously (**Supplemental table 6**)^42^. K-12 transconjugants carrying the urinary plasmids were able to grow on double chloramphenicol and tetracycline LB plates. For DNA extraction, bacteria were grown as liquid cultures in LB (with chloramphenicol and tetracycline as selection markers) at 37°C in a shaking incubator for 12 hours. Whole genome DNA was extracted via the Qiagen DNeasy UltraClean Microbial Kit using the standard manufacturer protocol. Plasmid DNA was extracted via the Qiagen Large-Construct Kit using the standard manufacturer protocol. A Qubit fluorometer was used to quantify the DNA concentration in the obtained genomic and plasmid DNA preparations. The Nextera XT DNA library preparation kit was used to make DNA libraries, which were sequenced on the Illumina MiSeq platform using the MiSeq Reagent Kit v2 (500-cycles) in Loyola University Chicago’s Genomics Facility (Maywood, IL, United States) Raw sequence reads for transconjugant K-12 respectively carrying pU1223 (SRR15011177) and pU1284 (SRR15011176) were submitted to GenBank. The raw sequence reads from the whole genome extraction were assembled using plasmidspades.py of SPAdes v3.12 with k values of 55,77,99,127 and the only-assembler parameter^43^. Assemblies were renamed via a bash script and contigs less than 500 bp were removed via Bioawk. Plasmidic assemblies were queried against the nr/nt database via NCBI’s web BLAST and hits were binned as either *E. coli* plasmid or chromosome^40,41^. A curated plasmidic assembly was made using only contigs hits to plasmid sequences by pruning contigs with homology to chromosomal entries in the nr/nt database.

NCBI Local BLAST v2.11.0 was used to query conjugation genes, as described above, in the plasmidic assemblies for plasmids pU1223 and pU1284^40,41^. pU0928 and pU1284 were annotated via Prokka v1.13.5 and the location of F-type and P-type conjugation genes was identified using the genome viewer function of Geneious Prime v2021.1prokk^44,45^. Predicted genes in these loci were queried against the NCBI nr/nt collection via BLAST^40,41^. Conjugation genes in pU1223 and pU1284 not included in our initial query list were concatenated into a new query list and all 65 urinary *E. coli* draft genomes were queried for these additional genes. Hits with a query coverage over 85% and sequence identity over 90% were recorded as present. We designated these parameters to have a high homology threshold while accounting for reported variable regions in conjugation genes^20^. The proportion of all genes queried, including those in the initial scan and those newly identified in pU1223 and pU1284, was calculated for all urinary *E. coli* genomes.

## Results

Sixty-five urinary *E. coli* draft genomes were queried by PlasmidSPAdes to identify plasmid *inc* genes (**Supplemental Table 2**). These 65 strains were isolated by our team via catheterization and thus are members of the bladder microbiota, which differs from the urethral microbiota^46^ that is also present in voided urine samples. For each genome sequence, we also have symptom data from the participant. To facilitate plasmid group analyzes, we binned isolates with *incF* genes into the IncF group, isolates with *col* genes but no *incF* genes in the Col group, and isolates with *inc* genes that were neither *incF* nor *col* into the Inc-various group. We performed an initial query utilizing F-type and P-type genes on 65 urinary *E. coli* draft genomes (genes queried are listed in **Supplemental Table 3 and 4)**^16,20^. Genes previously associated with F plasmid transfer, most notably *tra* genes, were in relatively high proportion in draft genomes with F-type genes (**Figure 1**). To validate plasmid conjugation in these isolates, we respectively chose two urinary *E. coli* isolate predicted to have a F-type system (UMB1223) and P-type system (UMB1284)^16^. UMB1223 and UMB1284 could grow on tetracycline plates and their tetracycline resistance gene was localized to a putative plasmid sequence within the genome assembly (**Supplemental Table 5**). Our initial gene query utilizing reference conjugation genes indicated that UMB1223 had 23 *tra* genes in a putative plasmid sequence, while UMB1284 had five *virB* genes in addition to two *tra* genes (**Supplemental Table 6**). To confirm conjugation, the genetic content in the MG1655 transconjugants was extracted, sequenced, and submitted to GenBank (K-12 with plasmid from UMB1223 as SRR15011177, K-12 with plasmid from UMB1284 as SRR15011176). The raw sequence reads were assembled into plasmid contigs (pU1223 and pU1284, respectively), and analyzed. We verified the presence of a tetracycline resistance cassette, *incF* loci, and conjugation genes like those present in the original (donor) urinary isolates (compare **Table 1** to **Supplemental Table 6**).

**Figure 1.**
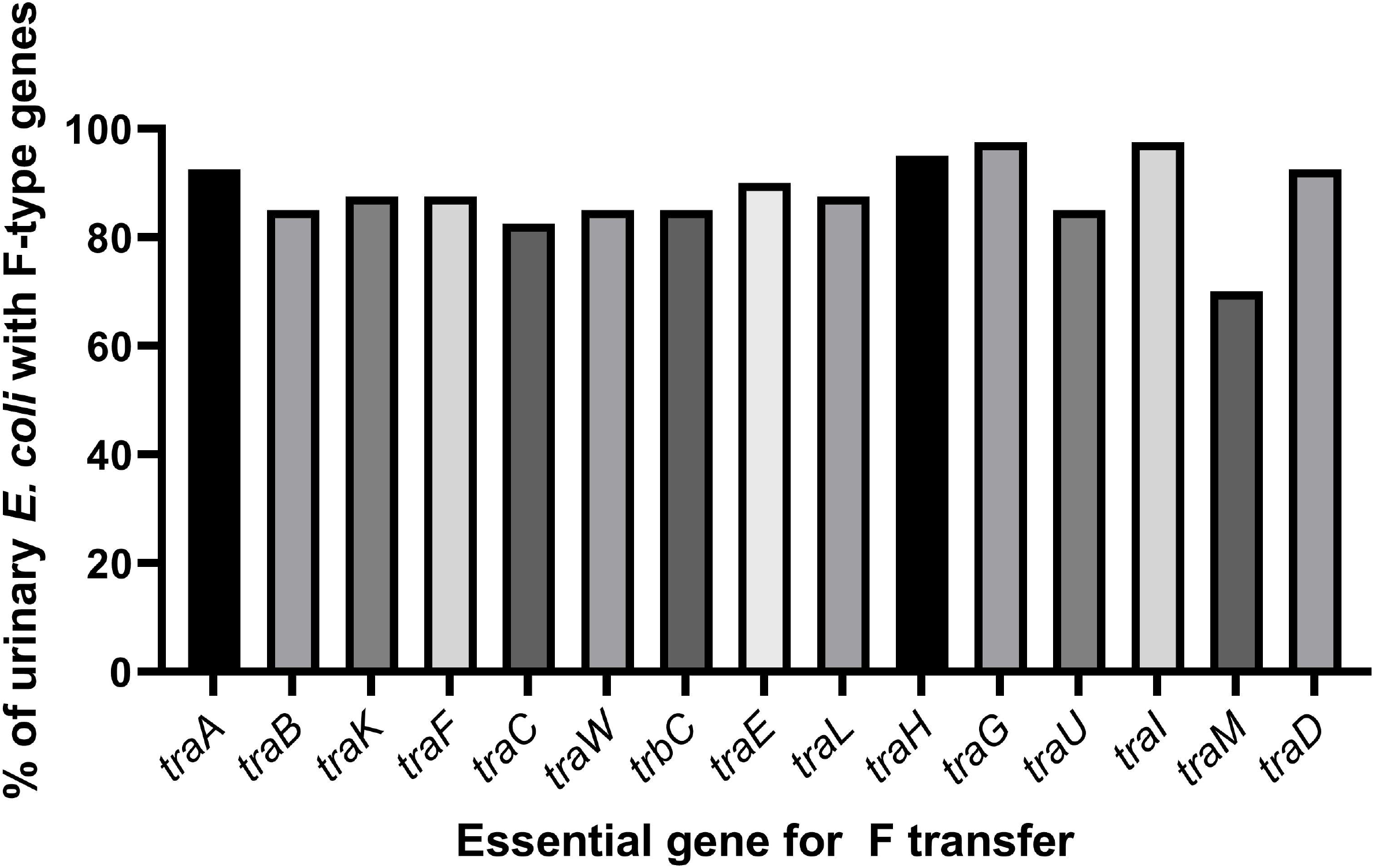
F transfer genes in sequences with evidence of an F-type system. Transfer genes were identified in urinary *E. coli* genomes. Genomes with evidence of an F-type system were further analyzed for the presence of transfer genes required for conjugation and could thus most likely succeed in conjugating their plasmid to *E. coli* K-12.

**Table 1.** Overview of urinary plasmids sequenced from *E. coli* K-12 transconjugants.

| Background Strain | Urinary *E. coli* plasmid donor | Urinary plasmid replicon | Tc resistance gene in plasmid | Conjugation genes identified<sup>442</sup><br><sup>443</sup> |
| --- | --- | --- | --- | --- |
| MG1655 | UMB1223 | col156, incFIA, incFIB, incFII | Yes | F-type |
| MG1655 | UMB1284 | incFIA, incFII, incX | Yes | P-type, F-type |

We reviewed the contigs in pU1223 and pU1284 for the presence of conjugation loci, including genes we may not have identified in our initial query using conjugation gene references (**Supplemental Table 3 and 4**). The conjugation locus in pU1223 consisted of 39 open reading frames (ORF) that included the transfer genes we had profiled in the parent isolate UMB1223; it also included variants of transfer genes not initially identified, two transposons, and four hypothetical genes (**Figure 2, Supplemental Table 7**). The transfer locus was organized in the prototypical manner of a F-type transfer operon, with *finO* upstream and both *traJ* and *traM* downstream of the rest of the *tra* genes, plus the presence of all *tra* genes essential for conjugative transfer^20,21^. The operon contained 17 non-essential transfer genes, in addition to 2 transposon sequences mid-operon, plus four hypothetical (HP) coding regions flanked by transfer genes*. HP_1* was 109 amino acids long and was not homologous to any entries in the NCBI database or known protein domains. *HP_2* had some homology to the neighboring *traC,* while *HP_3* and *HP_4* had homology to the neighboring *traJ.* Seven of the transfer genes identified in pU1223 were variants of the reference sequences we initially used to profile and were not initially identified because they failed to meet our threshold relative to the queries used; all these variants had high homology to transfer genes in the GenBank (**Supplemental Table 7**).

**Figure 2.**
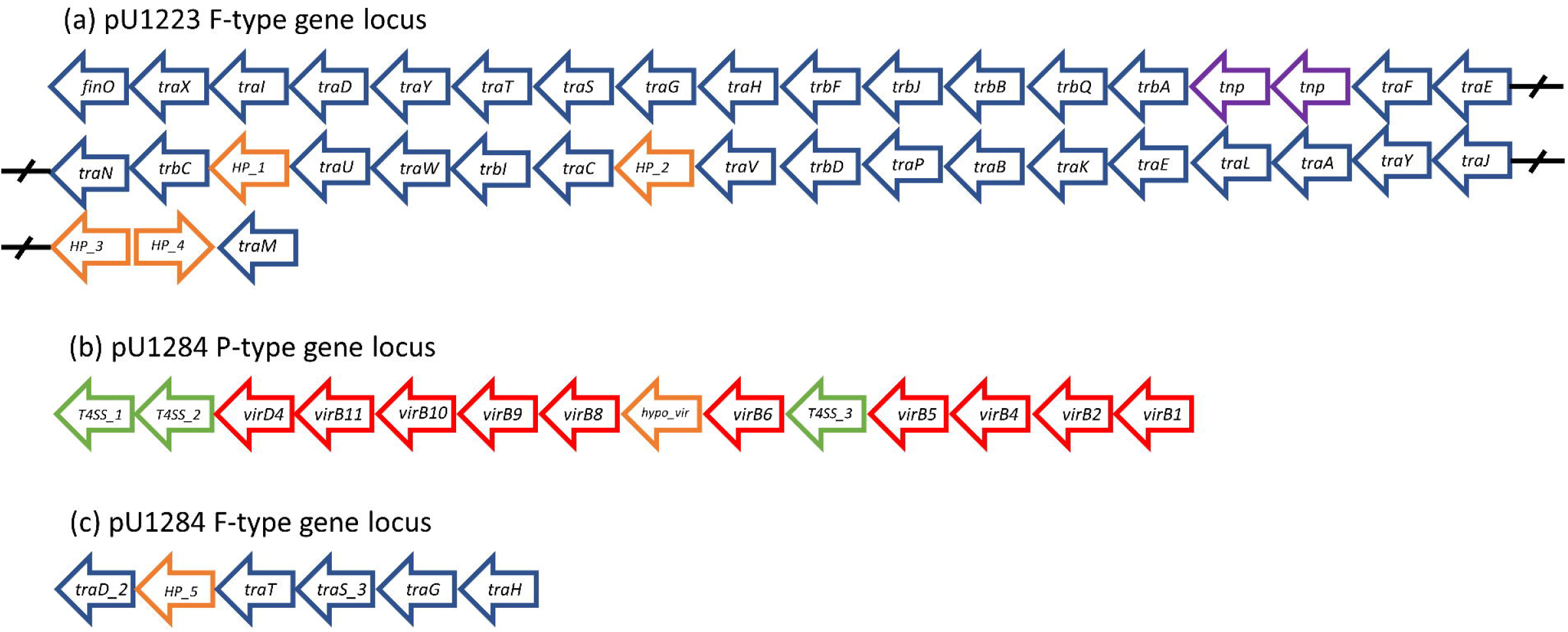
Conjugation gene operons in the plasmids conjugated to a *E. coli* K-12 strain from UMB1223 and UMB1284. Plasmidic contigs were assembled from the raw sequencing reads of *E. coli* K-12 strain MG1655 carrying a plasmid from either UMB1223 or UMB1284. Contigs containing hits to conjugation genes were identified. (a) pU1223 had a 39 gene operon containing *tra* and *trb* genes, in addition to two transposons (Tnsp). (b) pU1284 had an operon containing *vir* genes in one contig, including three unknown genes with homology to a T4SS and a hypothetical gene. In addition. pU1284 had a locus of *tra* genes in another contig, including a hypothetical gene.

pU1284 contained the P-type *virB-virD4* operon in one contig and a locus with F-type genes in another contig (**Figure 2**). Except for *virB3* and *virB7,* pU1284 had all VirB-VirD4 genes in the same locus with a prototypical sequential organization (**Supplemental Table 8**). Some of the *vir* components missed in our initial profile were present in the operon as a variant of our initial query sequence. We also identified three novel genes with T4SS homology upstream of *virD4,* between *virB6* and *virB5.* These T4SS ORFs had high sequence identity and query coverage to entries associated with conjugative and T4SS function in GenBank (**Supplemental Table 8**). T4SS_1 had a conjugal protein domain (PRK13893, pfam07424) like that in TrbM. T4SS_2 had protein domains of the Cag pathogenicity island proteins (pfam13117). T4SS_3 had the TrbL protein domain associated with conjugal proteins, in addition to a VirB6-like domain and conjugal protein domain like that in TrbL (pfam04610, COG3704, PRK13875). A hypothetical gene (hypo_*vir*) was also found flanked by *virB8* and *virB6,* where the missing *virB7* is normally located. In addition to the *virB-virD4* operon, pU1284 had a locus in another contig with transfer genes including *traD, traT, traS, traG, traH,* and a hypothetical ORF (*HP_5*) (**Supplemental Table 8**). *HP_5* had no distinct homology to entries in the NCBI database but was predicted to have multiple transmembrane domains.

We profiled all the urinary *E. coli* genomes using the newly identified ORFs in the putative conjugation regions from pU1223 and pU1284 (**Figure 3, Supplemental Table 9**). We estimated that plasmids are present in 84.6% (N=55/65) of these urinary *E. coli* isolates; most draft genomes had *incF* genes (81.8%, N=45/55). F-type conjugation genes were present in 72.7% of plasmids (N=40/55), all but one had *incF* genes (**Supplemental Table 9**). No conjugation genes were predicted in the genomes from the Col group (**Supplemental Table 9**). One urinary *E. coli* draft genome from the Inc-various group had 28 *tra* genes (**Supplemental Table 9**). Urinary *E. coli* with no *inc* gene hits did not have F-type or P-type conjugations genes (**Supplemental Table 9**). Nearly all F-type genes queried were present in draft genomes from the IncF group, with a mean gene hit count of 27.1 genes per genome (**Figure 3**). Multiple sequence variants were identified for *traJ, traS,* and *traY.* The genes *trbP, trwB, trwC,* and *trwM* were not found in any of the urinary *E. coli* genomes (**Supplemental Table 9**).

**Figure 3.**
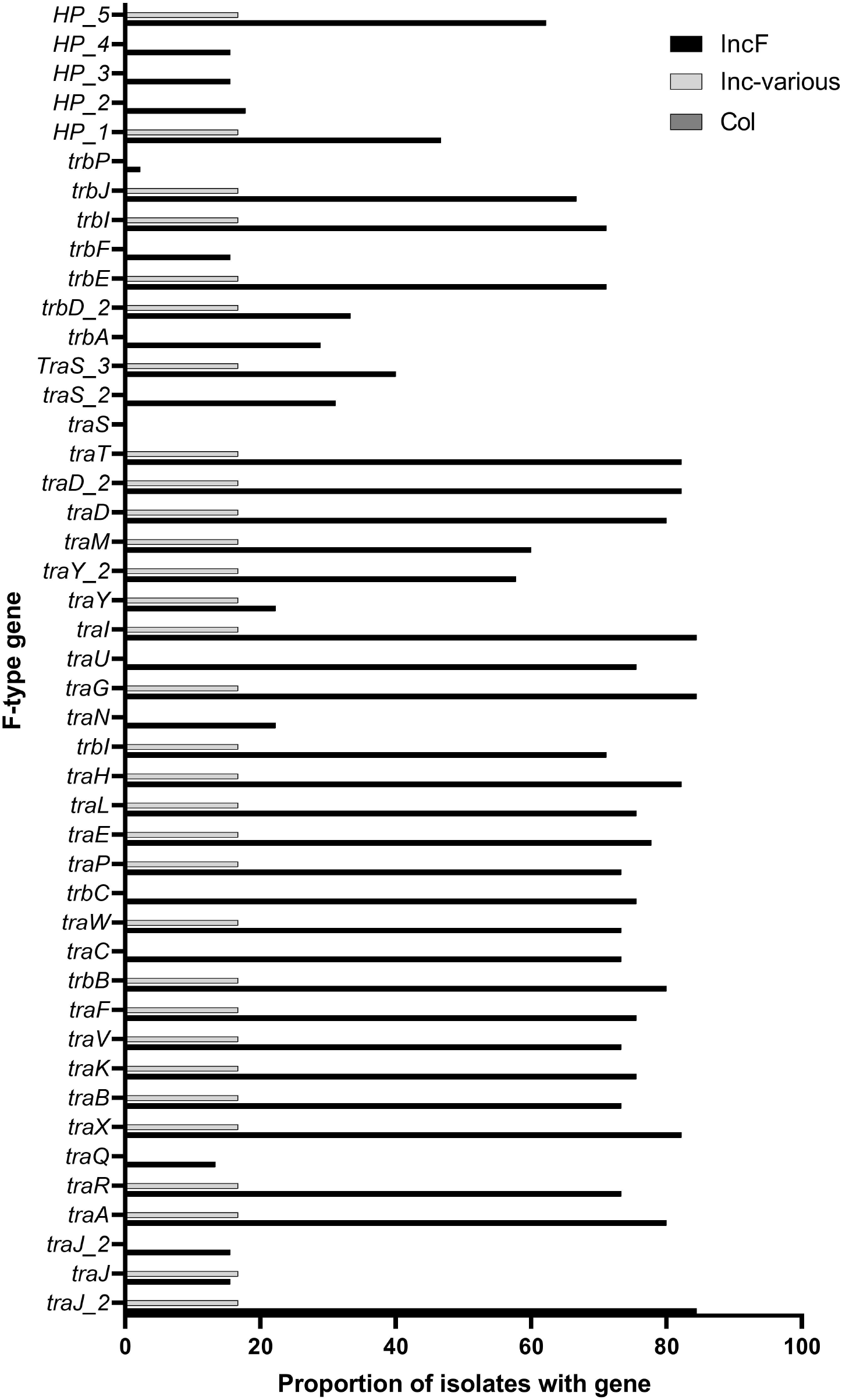
Proportion of *tra* genes in urinary *E. coli* plasmid group. The urinary *E. coli* genomes were scanned for all F-type genes, including variants and newly identified genes in pU1223 and pU1284. Urinary *E. coli* isolates with *incF* loci had a higher proportion than other plasmid groups in nearly every gene tested.

Type-P genes, consisting of the VirB-VirD4 system, were present in 18.2% (N=10/55) of the urinary *E. coli* isolates with putative plasmids (**Table 2**). Five genomes (UMB0931, UMB1284, UMB1727, UMB6721, UMB7431) were predicted to encode *vir* genes like those in pU1284, encompassing a near complete VirB-VirD4 operon. Urinary *E. coli* genomes with a near complete VirB-VirD4 operon (including pU1284) had the gene *incFII.* Except for UMB1348, draft genomes with a small number of P-type genes did not have the gene *incFII* but rather *incFI* (UMB6655, UMB6890) or *incX1* (UMB1228) (**Table 2**). Six urinary *E. coli* genomes (UMB0931, UMB1284, UMB1727, UMB6721, UMB7431, UMB6653) were predicted to encode the three novel T4SS-like ORFs.

**Table 2.**
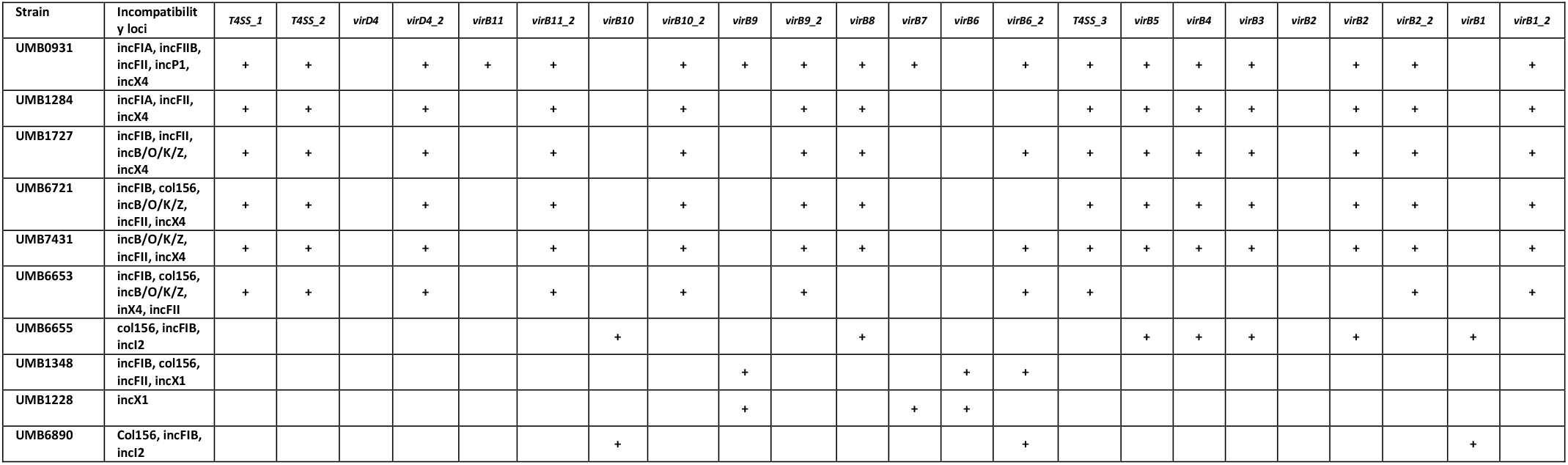
Urinary *E. coli* isolates with VirB-VirD4 genes.

## Discussion

Urinary *E. coli* are associated with UTI^1–4,47,48^. In urinary isolates of *E. coli,* virulence factors and antibiotic resistance cassettes are often retained and transmitted by conjugal plasmids^10,49^. F plasmids have been reported previously as present in the urinary tract, both in *E. coli* strains isolated from patients with UTI diagnoses and in an F-like plasmid hybrid in the UPEC strain UTI89^6,17,18^. Here, we have examined a representative set of urinary *E. coli* genomes from the bladders of adult women with and without UTI symptoms, providing evidence and investigation of the plasmid conjugation genes.

Conjugation components have been considered as targets to decrease the spread of plasmid-borne traits in the clinical setting^26,32^. For this strategy to be viable, we must understand the conjugation systems in the niche of interest. The urinary plasmid pU1223 had the F-type conjugation system, the most common conjugation system in the collection of urinary *E. coli* examined here (**Figure 2, Supplemental Table 7, Supplemental Table 8**). The conjugation operon in pU1223 had all genes deemed “essential” in an F-type system, in addition to 17 accessory genes; however, not all of the other urinary *E. coli* draft genomes followed this pattern^16,20^. While genes such as *traA, traH, traG, traI,* and *traD* were detected in more than 90% of draft genomes with F-type genes, there was no single gene present in all these genomes (**Figure 1, Figure 3**). We could infer that even if we targeted one of these high proportion components, a minority of urinary *E. coli* could potentially be unaffected.

Plasmid pU1284 had two conjugation loci, one a *virB-virD4* operon and another an F-type transfer locus (**Figure 2, Supplemental Table 8**). We identified VirB-VirD4 genes in ten urinary *E. coli* draft genomes (**Table 2, Supplemental Table 9**). Broadly, these urinary *E. coli* isolates with P-type genes fell into two groups. Plasmids with the first group had a near complete VirB-VirD4 operon similar to pU1284; all of these urinary *E. coli* genomes had the *incFII* gene. The second group had VirB-VirD4 genes that differed from those in pU1284; all but one lacked the *incFII* gene (**Table 2**). Urinary *E. coli* plasmids with P-type genes may represent a conjugative plasmid subpopulation; within this subpopulation, there may be further subgroups (i.e., those like pU1284 and those unlike pU1284)^14,38,50^. The presence of a minority of plasmids with a P-type conjugation system presents a troublesome scenario for targeting conjugation in this niche^14^. If we employ antibiotics against F-type genes, potentially we indirectly favor the propagation of plasmids with the P-type system^32,38^. Antibiotic use may have to be weighed with the risk that it will favor a subpopulation of organisms and potentially change the plasmid dynamics of the niche^38,51^.

A further layer to the diversity of conjugation systems in urinary *E. coli* is that pU1284 not only had a P-type operon, but also an F-type locus with the F-type conjugation genes *traD, traG,* and *traH* involved in the relaxosome, mating pair stabilization, and pilus retraction, respectively (**Figure 2, Supplemental table 8**). The F-type locus also had *traT* and *traS,* both involved in exclusion of invading plasmids in surface and entry exclusion, respectively^24,52^. In addition to UMB1284, five urinary *E. coli* (UMB0931, UMB1284, UMB1727, UMB6721, UMB6653, and UMB7431) had a VirB-VirD4 system and different combinations of F-type transfer genes (**Table 2, Supplemental Table 9**). These combinations of P-type and F-type genes could be linked to different environmental and evolutionary histories in the urinary tract^14,16,53^. This mosaic structure also highlights how urinary *E. coli* plasmids, such as F plasmids, can be genetically promiscuous^14^. Even if we target a conjugation system, this could be neutralized by the re-shuffling genetic content in these plasmids to generate mosaic conjugation loci.

An important factor to consider is that our urinary *E. coli* isolates were collected from the bladders of different participants^5^. Potentially, the diversity of conjugation genes in the urinary tract of a single person is much lower^54^. Notwithstanding, we postulate that whole genome sequencing and conjugation gene profiling would be key to the use of antibiotics that target conjugation components, especially in combination with traditional antibiotics^32^. Plasmid genetic variation in patients presents its own challenges, however, as targeting a single conjugation component may only be effective in a subset of cases. Overall, our evidence supports the concept that conjugation systems are common in urinary *E. coli,* especially in genomes with *incF* genes, and that these genes are diverse in number and composition^10^. The F-type system is the most common, but a minority of urinary *E. coli* had evidence of a P-type system^16^. Furthermore, urinary *E. coli* like UMB1284 appear to have supplemented their P-type system with some F-type genes, including a plasmid exclusion gene.

Conjugation components have been proposed as targets to limit the spread of plasmid-borne traits, such as antibiotic resistance^26,32^. Targeting of a specific conjugation component should be considered in the context of (1) how frequent is that component in the plasmid population, (2) how vital that component is to the conjugation system, and (3) whether another conjugation system in the population could be unscathed. Modularity and redundance is inherent to conjugation systems in bacteria and targeting isolated components could be ineffective or worse negatively affect the patient^10,14,16^. For example, by deploying antibiotics that target the F-type system, we may inadvertently increase the frequency of P-type conjugation, which could alter plasmid and microbiome dynamics^32^. The presence of different conjugation systems, frequency of individual conjugation genes, and combinations of these genes require further attention, as they may be key to plasmid exchange in the urinary tract. Conjugation systems in clinical isolates must be well characterized as part of our efforts to restrict the spread of plasmid-borne antibiotic resistance.

## ACKNOWLEDGMENTS

We thank Tanya Sysoeva (University of Alabama – Huntsville) for her critical review of our manuscript.

## Supplemental Information

Supplemental Table 1. List of urinary *E. coli* isolates used in this study

Supplemental Table 2. Inc loci in urinary *E. coli*

Supplemental Table 3. Transfer gene function and repository access

Supplemental Table 4. VirB-VirD4 and transfer gene homologues profiled in urinary *E. coli*

Supplemental Table 5. Growth of urinary *E. coli* isolates on antibiotic plates

Supplemental Table 6. Conjugation of urinary *E. coli* plasmids

Supplemental Table 7. Conjugation genes identified in pU1223

Supplemental Table 8. Conjugation genes identified in pU1284

Supplemental Table 9. Conjugation gene hits in urinary *E. coli* genomes

## REFERENCES

1. Foxman B. Urinary tract infection syndromes. Occurrence, recurrence, bacteriology, risk factors, and disease burden. Infect Dis Clin North Am. 2014;28(1):1–13. doi:10.1016/j.idc.2013.09.003

2. Flores-Mireles AL, Walker JN, Caparon M, Hultgren SJ. Urinary tract infections: Epidemiology, mechanisms of infection and treatment options. Nat Rev Microbiol. 2015;13(5):269–284. doi:10.1038/nrmicro3432

3. Ronald A. The etiology of urinary tract infection: Traditional and emerging pathogens. Am J Med. 2002;113(1 SUPPL. 1):14–19. doi:10.1016/S0002-9343(02)01055-0

4. Mao BH, Chang YF, Scaria J, et al. Identification of Escherichia coli genes associated with urinary tract infections. J Clin Microbiol. 2012;50(2):449–456. doi:10.1128/JCM.00640-11

5. Andrea Garretto, Taylor Miller-Ensminger, Adriana Ene, Zubia Merchant, Aashaka Shah, Athina Gerodias, Anthony Biancofiori, Stacey Canchola, Stephanie Canchola, Emanuel Castillo, Tasnim Chowdhury, Nikita Gandhi, Sarah Hamilton, Kyla Hatton, Syed Hyder, Kot AJW and CP. Genomic Survey of E. coli from the Bladders of Women with and without Lower Urinary Tract Symptoms. Front Microbiol Sect Infect Dis. 2020.

6. Chen SL, Hung C-S, Xu J, et al. Identification of genes subject to positive selection in uropathogenic strains of Escherichia coli: a comparative genomics approach. Proc Natl Acad Sci U S A. 2006;103(15):5977–5982. doi:10.1073/pnas.0600938103

7. Larsen P, Dynesen P, Nielsen KL, Frimodt-Møller N. Faecal Escherichia coli from patients with E. coli urinary tract infection and healthy controls who have never had a urinary tract infection. J Med Microbiol. 2014;63(4):582–589. doi:10.1099/jmm.0.068783-0

8. Koraimann G. Spread and Persistence of Virulence and Antibiotic Resistance Genes: A Ride on the F Plasmid Conjugation Module. EcoSal Plus. 2018;8(1). doi:10.1128/ecosalplus.esp-0003-2018

9. Steinig EJ, Duchene S, Robinson DA, et al. Evolution and global transmission of a multidrugresistant, community-associated methicillin-resistant staphylococcus aureus lineage from the Indian subcontinent. MBio. 2019;10(6). doi:10.1128/mBio.01105-19

10. Stephens C, Arismendi T, Wright M, et al. F Plasmids Are the Major Carriers of Antibiotic Resistance Genes in Human-Associated Commensal Escherichia coli. mSphere. 2020;5(4). doi:10.1128/msphere.00709-20

11. Mutai WC, Waiyaki PG, Kariuki S, Muigai AWT. Plasmid profiling and incompatibility grouping of multidrug resistant Salmonella enterica serovar Typhi isolates in Nairobi, Kenya. BMC Res Notes. 2019;12(1):1–6. doi:10.1186/s13104-019-4468-9

12. Arredondo-Alonso S, Willems RJ, van Schaik W, Schürch AC. On the (im)possibility of reconstructing plasmids from whole-genome short-read sequencing data. Microb Genomics. 2017;3(10). doi:10.1099/mgen.0.000128

13. Carattoli A, Zankari E, Garciá-Fernández A, et al. In Silico detection and typing of plasmids using plasmidfinder and plasmid multilocus sequence typing. Antimicrob Agents Chemother. 2014;58(7):3895–3903. doi:10.1128/AAC.02412-14

14. Osborn AM, da Silva Tatley FM, Steyn LM, Pickup RW, Saunders JR. Mosaic plasmids and mosaic replicons: Evolutionary lessons from the analysis of genetic diversity in IncFII-related replicons. Microbiology. 2000;146(9):2267–2275. doi:10.1099/00221287-146-9-2267

15. Yang QE, Sun J, Li L, et al. IncF plasmid diversity in multi-drug resistant Escherichia coli strains from animals in China. Front Microbiol. 2015;6(SEP). doi:10.3389/fmicb.2015.00964

16. Christie PJ. The Mosaic Type IV Secretion Systems. EcoSal Plus. 2016;7(1). doi:10.1128/ecosalplus.esp-0020-2015

17. Brolund A, Franzén O, Melefors Ö, Tegmark-Wisell K, Sandegren L. Plasmidome-Analysis of ESBL-Producing Escherichia coli Using Conventional Typing and High-Throughput Sequencing. PLoS One. 2013;8(6). doi:10.1371/journal.pone.0065793

18. Cusumano CK, Hung CS, Chen SL, Hultgren SJ. Virulence plasmid harbored by uropathogenic Escherichia coli functions in acute stages of pathogenesis. Infect Immun. 2010;78(4):1457–1467. doi:10.1128/IAI.01260-09

19. Smillie C, Garcillan-Barcia MP, Francia M V., Rocha EPC, de la Cruz F. Mobility of Plasmids. Microbiol Mol Biol Rev. 2010;74(3):434–452. doi:10.1128/mmbr.00020-10

20. Bragagnolo N, Rodriguez C, Samari-Kermani N, et al. Protein dynamics in f-like bacterial conjugation. Biomedicines. 2020;8(9). doi:10.3390/BIOMEDICINES8090362

21. Fernandez-Lopez R, de Toro M, Moncalian G, Garcillan-Barcia MP, de la Cruz F. Comparative genomics of the conjugation region of F-like plasmids: Five shades of F. Front Mol Biosci. 2016;3(NOV). doi:10.3389/fmolb.2016.00071

22. Lawley TD, Klimke WA, Gubbins MJ, Frost LS. F factor conjugation is a true type IV secretion system. FEMS Microbiol Lett. 2003;224(1):1–15. doi:10.1016/S0378-1097(03)00430-0

23. Virolle C, Goldlust K, Djermoun S, Bigot S, Lesterlin C. Plasmid transfer by conjugation in gramnegative bacteria: From the cellular to the community level. Genes (Basel). 2020;11(11):1–33. doi:10.3390/genes11111239

24. Achtman M, Kennedy N, Skurray R. Cell-cell interactions in conjugating Escherichia coli: Role of traT protein in surface exclusion. Proc Natl Acad Sci U S A. 1977;74(11):5104–5108. doi:10.1073/pnas.74.11.5104

25. Alt-Mörbe J, Stryker JL, Fuqua C, Li PL, Farrand SK, Winans SC. The conjugal transfer system of Agrobacterium tumefaciens octopine-type Ti plasmids is closely related to the transfer system of an IncP plasmid and distantly related to Ti plasmid vir genes. J Bacteriol. 1996;178(14):4248–4257. doi:10.1128/jb.178.14.4248-4257.1996

26. Cabezón E, de la Cruz F, Arechaga I. Conjugation inhibitors and their potential use to prevent dissemination of antibiotic resistance genes in bacteria. Front Microbiol. 2017;8(NOV). doi:10.3389/fmicb.2017.02329

27. Fronzes R, Schäfer E, Wang L, Saibil HR, Orlova E V., Waksman G. Structure of a type IV secretion system core complex. Science (80-). 2009;323(5911):266–268. doi:10.1126/science.1166101

28. Chandran V, Fronzes R, Duquerroy S, Cronin N, Navaza J, Waksman G. Structure of the outer membrane complex of a type IV secretion system. Nature. 2009;462(7276):1011–1015. doi:10.1038/nature08588

29. Low HH, Gubellini F, Rivera-Calzada A, et al. Structure of a type IV secretion system. Nature. 2014;508(7497):550–553. doi:10.1038/nature13081

30. Fronzes R, Christie PJ, Waksman G. The structural biology of type IV secretion systems. Nat Rev Microbiol. 2009;7(10):703–714. doi:10.1038/nrmicro2218

31. Zatyka M, Thomas CM. Control of genes for conjugative transfer of plasmids and other mobile elements. FEMS Microbiol Rev. 1998;21(4):291–319. doi:10.1111/j.1574-6976.1998.tb00355.x

32. Graf FE, Palm M, Warringer J, Farewell A. Inhibiting conjugation as a tool in the fight against antibiotic resistance. Drug Dev Res. 2019;80(1):19–23. doi:10.1002/ddr.21457

33. Sayer JR, Walldén K, Pesnot T, et al. 2- and 3-substituted imidazo[1,2-a]pyrazines as inhibitors of bacterial type IV secretion. Bioorganic Med Chem. 2014;22(22):6459–6470. doi:10.1016/j.bmc.2014.09.036

34. Shaffer CL, Good JAD, Kumar S, et al. Peptidomimetic small molecules disrupt type IV secretion system activity in diverse bacterial pathogens. MBio. 2016;7(2). doi:10.1128/mBio.00221-16

35. Ripoll-Rozada J, García-Cazorla Y, Getino M, et al. Type IV traffic ATPase TrwD as molecular target to inhibit bacterial conjugation. Mol Microbiol. 2016;100(5):912–921. doi:10.1111/mmi.13359

36. Bikard D, Euler CW, Jiang W, et al. Exploiting CRISPR-cas nucleases to produce sequence-specific antimicrobials. Nat Biotechnol. 2014;32(11):1146–1150. doi:10.1038/nbt.3043

37. Garcillán-Barcia MP, Jurado P, González-Pérez B, Moncalián G, Fernández LA, De La Cruz F. Conjugative transfer can be inhibited by blocking relaxase activity within recipient cells with intrabodies. Mol Microbiol. 2007;63(2):404–416. doi:10.1111/j.1365-2958.2006.05523.x

38. Arredondo-Alonso S, Top J, McNally A, et al. Plasmids shaped the recent emergence of the major nosocomial pathogen Enterococcus faecium. MBio. 2020;11(1). doi:10.1128/mBio.03284-19

39. Jalasvuori M, Friman V-P, Nieminen A, Bamford JKH, Buckling A. Bacteriophage selection against a plasmid-encoded sex apparatus leads to the loss of antibiotic-resistance plasmids. Biol Lett. 2011;7(6):902–905. doi:10.1098/rsbl.2011.0384

40. Altschul SF, Gish W, Miller W, Myers EW, Lipman DJ. Basic local alignment search tool. J Mol Biol. 1990;215(3):403–410. doi:10.1016/S0022-2836(05)80360-2

41. Camacho C, Coulouris G, Avagyan V, et al. BLAST+: Architecture and applications. BMC Bioinformatics. 2009;10. doi:10.1186/1471-2105-10-421

42. Barrick J. General conjugation protocol. https://barricklab.org/twiki/bin/view/Lab/ProtocolsConjugation. Published 2020. Accessed March 9, 2021.

43. Antipov D, Hartwick N, Shen M, Raiko M, Lapidus A, Pevzner PA. plasmidSPAdes: assembling plasmids from whole genome sequencing data. Bioinformatics. 2016;32(22):btw493. doi:10.1093/bioinformatics/btw493

44. Seemann T. Prokka: rapid prokaryotic genome annotation. Bioinformatics. 2014;30(14):2068–2069. doi:10.1093/bioinformatics/btu153

45. Geneious. Geneious 2021.0.3. https://www.geneious.com. Published 2021.

46. YB C, B H, TT P, M A-A, ER M, AJ W. The Urethral Microbiota: A Missing Link in the Female Urinary Microbiota. J Urol. 2020;204(2):303–309. doi:10.1097/JU.0000000000000910

47. Yamamoto S, Tsukamoto T, Terai A, Kurazono H, Takeda Y, Yoshida O. Genetic evidence supporting the fecal-perineal-urethral hypothesis in cystitis caused by Escherichia coli. J Urol. 1997;157(3):1127–1129. http://www.ncbi.nlm.nih.gov/pubmed/9072556. Accessed August 10, 2019.

48. Alteri CJ, Mobley HLT. Metabolism and Fitness of Urinary Tract Pathogens. Microbiol Spectr. 2015;3(3). doi:10.1128/microbiolspec.MBP-0016-2015

49. Zagaglia C, Casalino M, Colonna B, Conti C, Calconi A, Nicoletti M. Virulence plasmids of enteroinvasive Escherichia coli and Shigella flexneri integrate into a specific site on the host chromosome: Integration greatly reduces expression of plasmid-carried virulence genes. Infect Immun. 1991;59(3):792–799. doi:10.1128/iai.59.3.792-799.1991

50. Wagner S, Lupolova N, Gally DL, Argyle SA. Convergence of plasmid architectures drives emergence of multi-drug resistance in a clonally diverse Escherichia coli population from a veterinary clinical care setting. Vet Microbiol. 2017;211:6–14. doi:10.1016/j.vetmic.2017.09.016

51. Tepekule B, Wiesch PA Zur, Kouyos RD, Bonhoeffer S. Quantifying the impact of treatment history on plasmid-mediated resistance evolution in human gut microbiota. Proc Natl Acad Sci U S A. 2019;116(46):23106–23116. doi:10.1073/pnas.1912188116

52. Sukupolvil S, David O’connor C.2 TraT Lipoprotein, a Plasmid-Specified Mediator of Interactions between Gram-Negative Bacteria and Their Environment.; 1990. https://www.ncbi.nlm.nih.gov/pmc/articles/PMC372785/pdf/microrev00039-0011.pdf. Accessed June 17, 2019.

53. Zienkiewicz M, Kern-Zdanowicz I, Gołębiewski M, et al. Mosaic structure of p1658/97, a 125-kilobase plasmid harboring an active amplicon with the extended-spectrum β-lactamase gene blaSHV-5. Antimicrob Agents Chemother. 2007;51(4):1164–1171. doi:10.1128/AAC.00772-06

54. Skyberg JA, Johnson TJ, Johnson JR, Clabots C, Logue CM, Nolan LK. Acquisition of Avian Pathogenic Escherichia coli Plasmids by a Commensal E. coli Isolate Enhances Its Abilities To Kill Chicken Embryos, Grow in Human Urine, and Colonize the Murine Kidney. Infect Immun. 2006;74(11):6287–6292. doi:10.1128/IAI.00363-06

