## Supplemental table 1-8 for "Profiling the plasmid conjugation potential of urinary *E. coli*"

### Supplemental Information

**Supplemental Table 1. List of urinary *E. coli* isolates used in this study**

| Strain | Taxonomy | Assembly | Level | WGS | BioSample |
|---|---|---|---|---|---|
| UMB0103 | *Escherichia coli* | GCA_003892645.1 | Contig | RRWT00000000 | SAMN09665164 |
| UMB0149 | *Escherichia coli* | GCA_003892555.1 | Contig | RRWS00000000 | SAMN09665165 |
| UMB0276 | *Escherichia coli* | GCA_003892545.1 | Contig | RRWR00000000 | SAMN09665166 |
| UMB0527 | *Escherichia coli* | GCA_003892535.1 | Contig | RRWQ00000000 | SAMN09665167 |
| UMB0731 | *Escherichia coli* | GCA_003892485.1 | Contig | RRWP00000000 | SAMN09665168 |
| UMB0906 | *Escherichia coli* | GCA_003886695.1 | Contig | RRWO00000000 | SAMN09665169 |
| UMB0923 | *Escherichia coli* | GCA_003892635.1 | Contig | RRWN00000000 | SAMN09665170 |
| UMB0928 | *Escherichia coli* | GCA_003892445.1 | Contig | RRWM00000000 | SAMN09665171 |
| UMB0931 | *Escherichia coli* | GCA_003886495.1 | Contig | RRWL00000000 | SAMN09665172 |
| UMB0933 | *Escherichia coli* | GCA_003886675.1 | Contig | RRWK00000000 | SAMN09665173 |
| UMB0934 | *Escherichia coli* | GCA_003892475.1 | Contig | RRWJ00000000 | SAMN09665174 |
| UMB0939 | *Escherichia coli* | GCA_003885295.1 | Contig | RRUR00000000 | SAMN09665218 |
| UMB0949 | *Escherichia coli* | GCA_003892435.1 | Contig | RRWI00000000 | SAMN09665175 |
| UMB1012 | *Escherichia coli* | GCA_003886455.1 | Contig | RRWH00000000 | SAMN09665176 |
| UMB1091 | *Escherichia coli* | GCA_003886445.1 | Contig | RRWG00000000 | SAMN09665177 |
| UMB1093 | *Escherichia coli* | GCA_003885215.1 | Contig | RRUQ00000000 | SAMN09665219 |
| UMB1160 | *Escherichia coli* | GCA_003892605.1 | Contig | RRWF00000000 | SAMN09665178 |
| UMB1162 | *Escherichia coli* | GCA_003892455.1 | Contig | RRWE00000000 | SAMN09665179 |
| UMB1180 | *Escherichia coli* | GCA_008726795.1 | Contig | VYWI00000000 | SAMN12797014 |
| UMB1193 | *Escherichia coli* | GCA_003892595.1 | Contig | RRWC00000000 | SAMN09665181 |
| UMB1195 | *Escherichia coli* | GCA_008726745.1 | Contig | VYWF00000000 | SAMN12797017 |
| UMB1202 | *Escherichia coli* | GCA_003886395.1 | Contig | RRWA00000000 | SAMN09665183 |
| UMB1220 | *Escherichia coli* | GCA_003886385.1 | Contig | RRVZ00000000 | SAMN09665184 |
| UMB1221 | *Escherichia coli* | GCA_003885055.1 | Scaffold | RRUG00000000 | SAMN10411422 |
| UMB1223 | *Escherichia coli* | GCA_003886375.1 | Contig | RRVY00000000 | SAMN09665185 |
| UMB1225 | *Escherichia coli* | GCA_008726695.1 | Contig | VYWD00000000 | SAMN12797019 |

| | | | | | |
|---|---|---|---|---|---|
| UMB1228 | *Escherichia coli* | GCA_003886655.1 | Contig | RRVW00000000 | SAMN09665187 |
| UMB1229 | *Escherichia coli* | GCA_003886345.1 | Contig | RRVV00000000 | SAMN09665188 |
| UMB1284 | *Escherichia coli* | GCA_003892355.1 | Contig | RRVU00000000 | SAMN09665189 |
| UMB1285 | *Escherichia coli* | GCA_003886635.1 | Contig | RRVT00000000 | SAMN09665190 |
| UMB1335 | *Escherichia coli* | GCA_003886615.1 | Contig | RRVS00000000 | SAMN09665191 |
| UMB1337 | *Escherichia coli* | GCA_003886325.1 | Contig | RRVR00000000 | SAMN09665192 |
| UMB1346 | *Escherichia coli* | GCA_003886295.1 | Contig | RRVQ00000000 | SAMN09665193 |
| UMB1347 | *Escherichia coli* | GCA_003886285.1 | Contig | RRVP00000000 | SAMN09665194 |
| UMB1348 | *Escherichia coli* | GCA_003886275.1 | Contig | RRVO00000000 | SAMN09665195 |
| UMB1354 | *Escherichia coli* | GCA_003886225.1 | Contig | RRVN00000000 | SAMN09665196 |
| UMB1356 | *Escherichia coli* | GCA_003886245.1 | Contig | RRVM00000000 | SAMN09665197 |
| UMB1358 | *Escherichia coli* | GCA_003886195.1 | Contig | RRVL00000000 | SAMN09665198 |
| UMB1359 | *Escherichia coli* | GCA_003886185.1 | Contig | RRVK00000000 | SAMN09665199 |
| UMB1360 | *Escherichia coli* | GCA_003886565.1 | Contig | RRVJ00000000 | SAMN09665200 |
| UMB1362 | *Escherichia coli* | GCA_003886175.1 | Contig | RRVI00000000 | SAMN09665201 |
| UMB1526 | *Escherichia coli* | GCA_003886105.1 | Contig | RRVH00000000 | SAMN09665202 |
| UMB1727 | *Escherichia coli* | GCA_003886135.1 | Contig | RRVG00000000 | SAMN09665203 |
| UMB2019 | *Escherichia coli* | GCA_003886115.1 | Contig | RRVF00000000 | SAMN09665204 |
| UMB2055 | *Escherichia coli* | GCA_003886095.1 | Contig | RRVE00000000 | SAMN09665205 |
| UMB2321 | *Escherichia coli* | GCA_003886555.1 | Contig | RRVD00000000 | SAMN09665206 |
| UMB2328 | *Escherichia coli* | GCA_003886545.1 | Contig | RRVC00000000 | SAMN09665207 |
| UMB3538 | *Escherichia coli* | GCA_003886535.1 | Contig | RRVB00000000 | SAMN09665208 |
| UMB3641 | *Escherichia coli* | GCA_003885305.1 | Contig | RRUO00000000 | SAMN09665221 |
| UMB3643 | *Escherichia coli* | GCA_003885095.1 | Scaffold | RRUF00000000 | SAMN10411421 |
| UMB4656 | *Escherichia coli* | GCA_003886515.1 | Contig | RRVA00000000 | SAMN09665209 |
| UMB4716 | *Escherichia coli* | GCA_003885995.1 | Contig | RRUN00000000 | SAMN09665222 |
| UMB4746 | *Escherichia coli* | GCA_003886045.1 | Contig | RRUZ00000000 | SAMN09665210 |
| UMB5337 | *Escherichia coli* | GCA_003886035.1 | Contig | RRUY00000000 | SAMN09665211 |
| UMB5814 | *Escherichia coli* | GCA_003886015.1 | Contig | RRUX00000000 | SAMN09665212 |
| UMB5924 | *Escherichia coli* | GCA_003886005.1 | Contig | RRUW00000000 | SAMN09665213 |

| | | | | | |
|---|---|---|---|---|---|
| UMB5978 | *Escherichia coli* | GCA_003885915.1 | Contig | RRUV00000000 | SAMN09665214 |
| UMB6454 | *Escherichia coli* | GCA_003885245.1 | Contig | RRUU00000000 | SAMN09665215 |
| UMB6611 | *Escherichia coli* | GCA_003885875.1 | Contig | RRUT00000000 | SAMN09665216 |
| UMB6653 | *Escherichia coli* | GCA_003885965.1 | Scaffold | RRUS00000000 | SAMN09665217 |
| UMB6655 | *Escherichia coli* | GCA_003885255.1 | Contig | RRUL00000000 | SAMN09665225 |
| UMB6713 | *Escherichia coli* | GCA_003885145.1 | Contig | RRUK00000000 | SAMN09665226 |
| UMB6721 | *Escherichia coli* | GCA_003885125.1 | Scaffold | RRUJ00000000 | SAMN09665227 |
| UMB6890 | *Escherichia coli* | GCA_003885035.1 | Contig | RRUI00000000 | SAMN09665228 |
| UMB7431 | *Escherichia coli* | GCA_003885225.1 | Scaffold | RRUH00000000 | SAMN09665229 |

### Supplemental Table 3. Transfer gene function and repository access

| ***tra* gene** | **UniProtKB** |
|---|---|
| Transcription regulation | |
| *finO* | P22707 |
| *traJ* | P06626 |
| *traR* | P41065 |
| ProPilin maturation | |
| *traA* | P04737 |
| *traQ* | P18033 |
| *traX* | P22709 |
| T4SS core proteins | |
| *traB* | P41067 |
| *traK* | P41066 |
| *traV* | P41069 |
| Pilus assembly/extension | |
| *traF* | P14497 |
| *trbB* | P18035 |
| *traC* | P18004 |
| *traW* | P18472 |
| *trbC* | P18473 |
| *traP* | P41068 |
| *traE* | P08322 |
| *traL* | P08321 |
| Pilus retraction | |
| *traH* | P15069 |
| *trbI* | P18006 |
| Mating pair stabilization | |
| *traN* | P24082 |
| *traG* | P33790 |
| *traU* | P18471 |
| Relaxase | |
| *traI* | P14565 |
| Relaxosome accessory | |
| *traY* | P06627 |
| *traM* | P10026 |
| *traD* | P09130 |
| Surface/entry exclusion | |
| *traT* | P13979 |
| *traS* | P09129 |

**Supplemental Table 4. VirB-VirD4 and transfer gene homologues profiled in urinary *E. coli***

| Conjugation gene | tra homologues | UniProtKB |
|---|---|---|
| *virB1* | | A0A0R6LN99 |
| *virB2* | *traA* | A0A4C4D6J9 |
| *virB3* | *traL* | A0A4C4D6J9 |
| *virB4* | *traC* | A0A0A1DZA0 |
| *virB5* | *traE* | G8GYF8 |
| *virB6* | *traG* | G1CCR5 |
| *virB7* | *traV* | B0ZDY1 |
| *virB8* | *traG* | G8GYF5 |
| *virB9* | *traK* | A0A0C5F896 |
| *virB10* | *traB* | A0A0R6LEV8 |
| *virB11* | | A0A0C5EWI7 |
| *virD2* | *traI* | A0A482M9K1 |
| *virD4* | *traD* | A0A381CEJ0 |
| *ptleE* | | A0A377MWU4 |
| *trbC* | *traA* | A0A1P8KHV0 |
| *trbE* | *traC* | Q05807 |
| *trbI* | *traB* | P18006 |
| *trbP* | *traX* | W6AWJ8 |
| *trwB* | *traD* | Q04230 |
| *trwC* | *traI* | Q47673 |
| *trwM* | | O50329 |

**Supplemental Table 5. Growth of urinary *E. coli* isolates on antibiotic plates**

| | | | LB | Amp | Cm | Kan | Spec | Tet |
|---|---|---|---|---|---|---|---|---|
| | | **Total** | **68** | **28** | **1** | **3** | **8** | **16** |
| **Strain** | **Inc group** | **# abx that it grew on** | **100** | **41.1765** | **1.47059** | **4.41176** | **11.7647** | **23.5294** |
| B | Control | 0 | Yes | No | No | No | No | No |
| C | Control | 0 | Yes | No | No | No | No | No |
| K-12 | Control | 0 | Yes | No | No | No | No | No |
| UMB0103 | IncF | 3 | Yes | Yes | Yes | No | No | Yes |
| UMB0149 | Inc-various | 0 | Yes | No | No | No | No | No |
| UMB0276 | No inc | 1 | Yes | Yes | No | No | No | No |
| UMB0527 | IncF | 0 | Yes | No | No | No | No | No |
| UMB0731 | Col | 0 | Yes | No | No | No | No | No |
| UMB0906 | IncF | 1 | Yes | Yes | No | No | No | No |
| UMB0923 | Inc-various | 0 | Yes | No | No | No | No | No |
| UMB0928 | IncF | 2 | Yes | Yes | No | No | No | Yes |
| UMB0931 | IncF | 2 | Yes | Yes | No | No | No | Yes |
| UMB0933 | IncF | 1 | Yes | No | No | No | Yes | No |
| UMB0934 | IncF | 2 | Yes | Yes | No | No | No | Yes |
| UMB0939 | Col | 2 | Yes | Yes | No | No | No | Yes |
| UMB0949 | IncF | 2 | Yes | Yes | No | No | No | Yes |
| UMB1012 | IncF | 1 | Yes | Yes | No | No | No | No |
| UMB1091 | IncF | 3 | Yes | Yes | No | No | Yes | Yes |
| UMB1093 | IncF | 1 | Yes | Yes | No | No | No | No |
| UMB1160 | IncF | 1 | Yes | No | No | No | Yes | No |
| UMB1162 | IncF | 1 | Yes | No | No | No | No | Yes |
| UMB1180 | Col | 0 | Yes | No | No | No | No | No |
| UMB1193 | IncF | 3 | Yes | Yes | No | No | Yes | Yes |
| UMB1195 | IncF | 0 | Yes | No | No | No | No | No |
| UMB1202 | IncF | 0 | Yes | No | No | No | No | No |
| UMB1220 | No inc | 0 | Yes | No | No | No | No | No |
| UMB1221 | IncF | 1 | Yes | No | No | No | No | Yes |
| UMB1223 | IncF | 2 | Yes | Yes | No | No | No | Yes |
| UMB1225 | No inc | 0 | Yes | No | No | No | No | No |
| UMB1228 | Inc-various | 0 | Yes | No | No | No | No | No |
| UMB1229 | IncF | 3 | Yes | Yes | No | No | Yes | Yes |
| UMB1284 | IncF | 4 | Yes | Yes | No | Yes | Yes | Yes |
| UMB1285 | IncF | 0 | Yes | No | No | No | No | No |

| | | | | | | | | |
|---|---|---|---|---|---|---|---|---|
| UMB1335 | IncF | 1 | Yes | Yes | No | No | No | No |
| UMB1337 | IncF | 1 | Yes | Yes | No | No | No | No |
| UMB1346 | IncF | 0 | Yes | No | No | No | No | No |
| UMB1347 | IncF | 0 | Yes | No | No | No | No | No |
| UMB1348 | IncF | 1 | Yes | Yes | No | No | No | No |
| UMB1354 | No inc | 0 | Yes | No | No | No | No | No |
| UMB1356 | No inc | 0 | Yes | No | No | No | No | No |
| UMB1358 | No inc | 0 | Yes | No | No | No | No | No |
| UMB1359 | No inc | 0 | Yes | No | No | No | No | No |
| UMB1360 | IncF | 1 | Yes | Yes | No | No | No | No |
| UMB1362 | IncFI | 2 | Yes | Yes | No | No | No | Yes |
| UMB1526 | No inc | 1 | Yes | Yes | No | No | No | No |
| UMB1727 | IncFII | 1 | Yes | Yes | No | No | No | No |
| UMB2019 | Col | 0 | Yes | No | No | No | No | No |
| UMB2055 | Inc-various | 0 | Yes | No | No | No | No | No |
| UMB2321 | Inc-various | 0 | Yes | No | No | No | No | No |
| UMB2328 | Inc-various | 0 | Yes | No | No | No | No | No |
| UMB3538 | IncF | 2 | Yes | Yes | No | Yes | No | No |
| UMB3641 | IncF | 2 | Yes | Yes | No | No | Yes | No |
| UMB3643 | Inc-various | 1 | Yes | Yes | No | No | No | No |
| UMB4656 | IncF | 1 | Yes | Yes | No | No | No | No |
| UMB4716 | IncF | 0 | Yes | No | No | No | No | No |
| UMB4746 | IncF | 0 | Yes | No | No | No | No | No |
| UMB5337 | No inc | 0 | Yes | No | No | No | No | No |
| UMB5814 | IncF | 0 | Yes | No | No | No | No | No |
| UMB5924 | No inc | 4 | Yes | Yes | No | Yes | Yes | Yes |
| UMB5978 | IncF | 0 | Yes | No | No | No | No | No |
| UMB6454 | IncF | 0 | Yes | No | No | No | No | No |
| UMB6471 | IncF | 0 | Yes | No | No | No | No | No |
| UMB6611 | IncF | 0 | Yes | No | No | No | No | No |
| UMB6655 | IncF | 0 | Yes | No | No | No | No | No |
| UMB6713 | IncF | 1 | Yes | Yes | No | No | No | No |
| UMB6721 | IncF | 1 | Yes | No | No | No | No | Yes |
| UMB6890 | IncF | 0 | Yes | No | No | No | No | No |
| UMB7431 | IncF | 0 | Yes | No | No | No | No | No |

**Supplemental Table 6. Conjugation of urinary *E. coli* plasmids**

| Isolate | Role in conjugation | Urinary plasmid replicon | Conjugation genes profiled prior to conjugation | Tc resistance predicted[1] | Growth on Tc plate | Growth on Cm plate | Growth on Tc Cm plate | Produces transconjugants on Tc Cm plates[2] |
|---|---|---|---|---|---|---|---|---|
| K-12 MG1655 | Plasmid recipient | None | None | No | No | Yes | No | Yes |
| UMB1223 | Plasmid donor | IncFA, IncFIB(AP001918), IncFII(pRSB107) | *finO, traA, traR, traX, traB, traK, traV, traF, trbB, traC, traW, trbC, traP, traE, traL, traH, trbI, traG, traU, traI, traM, traD, traT* | Yes | Yes | No | No | Yes |
| UMB1284 | Plasmid donor | IncFIA, IncFII, IncX4 | *traG, traT, virB2, virB3, virB4, virB5, virB8* | Yes | Yes | No | No | Yes |
| UMB0939 | Plasmid donor without conjugation system (negative control) | Col(MG828), ColRNAI | None | No | Yes | No | No | No |

[1] Contig with gene for Tc resistance has homology to plasmid

[2] Urinary *E. coli* conjugated with *E. coli* K-12 strain MG1655 with chloramphenicol resistance selection marker

**Supplemental Table 7. Conjugation genes identified in pU1223**

| Annotation ID | Gene | Query Cover | E value | Per. Ident | Acc. Len | Accession |
|---|---|---|---|---|---|---|
| GJKJIFEI_00008 | *finO* | 100% | 8.00E-132 | 100.00% | 188 | KUT64489.1 |
| GJKJIFEI_00009 | *traX* | 100% | 2.00E-177 | 99.60% | 248 | WP_053879992.1 |
| GJKJIFEI_00010 | *traI* | 100% | 0 | 99.94% | 1756 | WP_172694226.1 |
| GJKJIFEI_00011 | *traD* | 100% | 0 | 99.73% | 735 | EHI0361888.1 |
| GJKJIFEI_00012 | *traY* | 100% | 0 | 100.00% | 287 | PHN11796.1 |
| GJKJIFEI_00013 | *traT* | 100% | 0 | 100.00% | 293 | ANK07087.1 |
| GJKJIFEI_00014 | *TraS_2* | 100% | 6.00E-117 | 99.39% | 163 | CTS32472.1 |
| GJKJIFEI_00015 | *traG* | 100% | 0 | 100.00% | 941 | HAM6569033.1 |
| GJKJIFEI_00016 | *traH* | 100% | 0 | 99.78% | 457 | WP_061089898.1 |
| GJKJIFEI_00017 | *trbF_2* | 100% | 2.00E-92 | 100.00% | 141 | WP_001348758.1 |
| GJKJIFEI_00018 | *trbJ_2* | 100% | 6.00E-61 | 98.95% | 95 | KUV34015.1 |
| GJKJIFEI_00019 | *trbB* | 100% | 2.00E-130 | 99.45% | 181 | WP_087507405.1 |
| GJKJIFEI_00020 | *traQ* | 100% | 7.00E-61 | 98.94% | 94 | HAJ2583838.1 |
| GJKJIFEI_00021 | *trbA_2* | 100% | 1.00E-68 | 100.00% | 127 | ESA77523.1 |
| GJKJIFEI_00022 | Transposase | 100% | 3.00E-120 | 99.40% | 167 | AVJ76634.1 |
| GJKJIFEI_00023 | Transposase | 100% | 0 | 99.69% | 326 | HAW2113186.1 |
| GJKJIFEI_00024 | *traF* | 100% | 0 | 100.00% | 257 | ARX27828.1 |
| GJKJIFEI_00025 | *trbE* | 100% | 9.00E-54 | 100.00% | 89 | WP_033554899.1 |
| GJKJIFEI_00026 | *traN* | 100% | 0 | 99.84% | 616 | HAJ2996344.1 |
| GJKJIFEI_00027 | *trbC* | 100% | 9.00E-156 | 100.00% | 216 | ARX27831.1 |
| GJKJIFEI_00028 | *HP_1* | 100% | 1.00E-67 | 100.00% | 109 | WP_139581483.1 |
| GJKJIFEI_00029 | *traU* | 100% | 0 | 99.70% | 330 | WP_021528459.1 |
| GJKJIFEI_00030 | *traW* | 100% | 2.00E-151 | 100.00% | 228 | RBW13522.1 |
| GJKJIFEI_00031 | *trbI* | 100% | 2.00E-88 | 99.22% | 133 | ADL14037.1 |
| GJKJIFEI_00032 | *traC* | 100% | 0 | 99.89% | 875 | WP_157352444.1 |
| GJKJIFEI_00033 | *HP_2* | 100% | 2.00E-15 | 100.00% | 48 | EHV46867.1 |
| GJKJIFEI_00034 | *traV* | 100% | 4.00E-120 | 99.42% | 171 | WP_097762797.1 |
| GJKJIFEI_00035 | *trbD_2* | 100% | 2.00E-80 | 100.00% | 127 | HAL3406214.1 |
| GJKJIFEI_00036 | *traP* | 100% | 5.00E-138 | 100.00% | 188 | WP_001617877.1 |
| GJKJIFEI_00037 | *traB* | 100% | 0 | 100.00% | 532 | WP_077758417.1 |

| | | | | | | |
|---|---|---|---|---|---|---|
| GJKJIFEI_00038 | *traK* | 100% | 4.00E-175 | 99.59% | 242 | WP_050008704.1 |
| GJKJIFEI_00039 | *traE* | 100% | 4.00E-136 | 99.47% | 188 | WP_032153574.1 |
| GJKJIFEI_00040 | *traL* | 100% | 4.00E-70 | 100.00% | 109 | WP_160372682.1 |
| GJKJIFEI_00041 | *traA* | 100% | 4.00E-77 | 99.17% | 120 | WP_069067339.1 |
| GJKJIFEI_00042 | *yraY_2* | 100% | 2.00E-45 | 100.00% | 81 | WP_049086340.1 |
| GJKJIFEI_00043 | *traJ_2* | 100% | 7.00E-49 | 100.00% | 83 | TFM56752.1 |
| GJKJIFEI_00044 | *HP_3* | 97% | 9.00E-19 | 97.67% | 43 | EGR70770.1 |
| GJKJIFEI_00045 | *HP_4* | 100% | 1.00E-32 | 100.00% | 59 | AKE87743.1 |
| GJKJIFEI_00046 | *traM* | 100% | 2.00E-86 | 99.21% | 127 | HAX8410798.1 |

**Supplemental Table 8. Conjugation genes identified in pU1284**

| Annotation ID | Gene | Query Cover | E value | Per. Ident | Acc. Len | Accession |
|---|---|---|---|---|---|---|
| OJDLIIBG_00019 | *T4SS_1* | 100% | 1.00E-65 | 99.01% | 101 | EFH9106740.1 |
| OJDLIIBG_00020 | *T4SS_2* | 100% | 3.00E-98 | 99.28% | 140 | OJM06430.1 |
| OJDLIIBG_00021 | *virD4_2* | 100% | 0 | 99.84% | 620 | WP_187440460.1 |
| OJDLIIBG_00022 | *virB11_2* | 100% | 0 | 99.71% | 342 | WP_089644540.1 |
| OJDLIIBG_00023 | *virB10_2* | 100% | 0 | 99.73% | 372 | EES3164312.1 |
| OJDLIIBG_00024 | *virB9* | 100% | 0 | 99.67% | 302 | WP_137486256.1 |
| OJDLIIBG_00025 | *virB8* | 100% | 5.00E-166 | 99.56% | 228 | WP_096190789.1 |
| OJDLIIBG_00026 | *hypo_vir* | 100% | 5.00E-35 | 100.00% | 67 | WP_000903502.1 |
| OJDLIIBG_00027 | *virB6_2* | 55% | 2.00E-10 | 84.85% | 397 | AQZ20251.1 |
| OJDLIIBG_00028 | *T4SS_3* | 100% | 0 | 100.00% | 364 | WP_125121539.1 |
| OJDLIIBG_00029 | *virB5* | 100% | 6.00E-174 | 99.58% | 238 | EFD0911387.1 |
| OJDLIIBG_00030 | *virB4* | 100% | 0 | 99.78% | 915 | EFB7338537.1 |
| OJDLIIBG_00031 | *virB2_2* | 100% | 1.00E-69 | 100.00% | 127 | ARH02006.1 |
| OJDLIIBG_00032 | *virB1_2* | 100% | 4.00E-150 | 99.51% | 206 | WP_032239609.1 |
| OJDLIIBG_00138 | *traD_2* | 99% | 1.00E-73 | 100.00% | 119 | EER1412495.1 |
| OJDLIIBG_00139 | *HP_5* | 100% | 0 | 99.59% | 245 | EFF9468201.1 |
| OJDLIIBG_00140 | *traT* | 100% | 1.00E-175 | 100.00% | 244 | AET14958.1 |
| OJDLIIBG_00141 | *traS_3* | 100% | 2.00E-112 | 98.11% | 159 | SRA12240.1 |

| | | | | | | |
|---|---|---|---|---|---|---|
| OJDLIIBG_00142 | *traG* | 100% | 0 | 99.89% | 940 | WP_097760672.1 |
| OJDLIIBG_00143 | *traH* | 100% | 0 | 100.00% | 332 | GDF19761.1 |
